# Bmal1 expression is minimal or absent in human and mouse cerebral microglia

**DOI:** 10.1101/2025.07.11.664407

**Authors:** Reza Rahimian, Sophie Simard, Anjali Chawla, Lei Zhu, Gustavo Turecki, Kai-Florian Storch, Naguib Mechawar

**Author notes:** **Corresponding authors:** Naguib Mechawar Ph.D., McGill Group for Suicide Studies, Douglas Mental Health Institute, Department of Psychiatry, McGill University, Kai-Florian Storch, Douglas Mental Health University Institute, McGill University, Montreal, QC H4H 1R3, Canada. Equal contribution (Reza Rahimian, Sophie Simard, Anjali Chawla).

## Abstract

Microglia orchestrate immunological responses in the brain and play an important role in maintaining homeostatic brain functions. Several studies have reported clock gene expression in microglia and the circadian rhythm they drive has been linked to the modulation of immune responses and neuronal functions. In the current study, complementary approaches, including immunofluorescence, multiplexed fluorescence *in situ* hybridization, and liquid chromatography-mass spectrometry proteomics of isolated CD11b^+^ microglia, were combined with publicly available transcriptomic and epigenomic datasets to investigate the expression of the core clock gene *BMAL1* in human post-mortem cortical and limbic areas as well as mouse brain. The majority of *BMAL1*-expressing cells were found to be neurons, with microglia representing a negligeable proportion. We also identified significantly lower chromatin accessibility or “openness” for *BMAL1* gene regulatory regions (such as promoters and enhancers) in microglia compared to neurons. These regulatory regions in microglia were enriched for ETS domain transcription factor (TF) binding sites. Together, this suggests a strong role of chromatin remodeling factors in suppressing *BMAL1* gene expression in microglia. Finally, while we observed a very low expression, BMAL1 TF motifs were accessible in open chromatin landscape of microglia, which may lead to downstream gene-regulatory effects upon binding, even if *BMAL1* expression is constitutively low. Overall, our results reveal low or absent expression of *BMAL1* in microglia and point towards potential epigenetic mechanisms regulating its expression in these cells.

## Main

Microglia as central nervous system (CNS) resident macrophages are instrumental in initiating the innate immune response against pathogens and toxins. In addition to developmental stage, brain region, and pathological state, different lines of evidence demonstrate that circadian time is a pivotal factor for microglial function in the healthy brain and in pathologies, as recently reviewed (Guzman-Ruiz et al., 2023; Jiao et al., 2024). Previous research has in fact indicated that microglia display exacerbated inflammatory responses depending on the time of day (Fonken et al., 2016; Guzman-Ruiz et al., 2023), but also express core clock genes rhythmically without the need for an immune stimulus (Nakazato et al., 2011). Rhythmic clock gene expression in isolated or cultured microglia has been indeed confirmed by others (Fonken et al., 2015; Hayashi et al., 2013; Nakazato et al., 2017; Nakazato et al., 2011; Ni et al., 2019; Wang et al., 2021). Among the clock genes shown to be expressed in microglia is the transcription factor Bmal1, which forms a dimer with Clock and binds to specific circadian promoter elements, so called E-boxes, to drive circadian gene expression of other clock components such as Period 1, 2, and 3, as well as Cryptochrome 1 and 2 genes (Dunlap, 1999; Schrader et al., 2024; Zhang & Kay, 2010). These clock genes form the negative limb of the core clock mechanism, which, upon their translation, move into the nucleus to act as inhibitors of their own transcription. Once their levels are low enough due to the sustained suppression of transcriptional activity, as well as their active degradation via the proteasomal pathway, inhibition is lifted, and the circadian gene expression cycle can start anew. In addition to this core transcriptional-translational feedback, there are other feedback loops contributing to the rhythm generation process, including one that involves the transcription factor Rev-Erb alpha which, as the Per and Cry genes, is driven by E-boxes but then, different from the Pers and Crys, acts as a transcriptional inhibitor at the Bmal1 locus, suppressing Bmal1’s expression rhythmically (Preitner et al., 2002). Reflective of its role as a core transcription factor in the clock mechanism, Bmal1 is essential for rhythm generation and thus, global *Bmal1^−/−^* mice are circadian arrhythmic throughout the body axis (Bunger et al., 2000).

During neuroinflammation, it has been shown that hippocampal microglia in rodents express high amounts of Bmal1 and pro-inflammatory cytokines during the light phase (Fonken et al., 2015). In addition, Bmal1 deficiency diminishes mRNA levels of canonical inflammatory markers such as TNF-α and IL-6 in response to LPS challenge *in vitro* (Wang et al., 2020). Furthermore, Wang et al. (2021) have previously demonstrated that microglia-specific disruption of the *Bmal1* gene improves long-term memory and protects mice from high fat diet-induced obesity. These and other studies typically examined Bmal1 expression in isolated or cultured microglia, but not in vivo. However, it is well established that culturing microglia can affect their phenotype, secretary profile, and gene expression profile in rodents (Mizrachi & Diamond, 2024) and in humans (Tewari et al., 2021). More importantly, comprehensive histological studies investigating microglial Bmal1 expression in different brain regions, in both humans and rodents, are lacking. To fill this knowledge gap, in this study, we used multiplexed fluorescence *in situ* hybridization (FISH), liquid chromatography-mass spectrometry (LC-MS) proteomics, as well as existing single-nucleus RNA sequencing (snRNA-sequencing), single-nucleus ATAC sequencing (snATAC-sequencing), and spatial transcriptomic (ST) datasets to better understand the phenotype of *BMAL1-*expressing cells in the human brain at the proteomic, transcriptomic, and epigenomic levels. We complemented these investigations with immunohistochemistry on mouse brain.

### BMAL1 is expressed in RBFOX3^+^ neurons but is lacking in P2RY12^+^ microglia in various regions of the human brain

While much of the literature on microglial clock function has been overwhelmingly focused on the cortex and the hippocampus (Guzman-Ruiz et al., 2023; Jiao et al., 2024), little is known about *BMAL1* microglial expression in other regions of the brain. Interestingly, a previous post-mortem human study found *BMAL1* as one of the top ranked rhythmic genes in the dorsolateral prefrontal cortex (dlPFC), anterior cingulate cortex (ACC), hippocampus, amygdala, nucleus accumbens, and cerebellum of the neurotypical brain (Li et al., 2013). However, the authors did not investigate cell-class specific *BMAL1* expression. To shed light on *BMAL1* expression according to cell class in the human brain, we first used multiplexed FISH (Advanced Cell Diagnostics RNAscope) for the visualization of *BMAL1* transcripts in cells expressing *P2RY12*, a purinergic receptor shown to be exclusively expressed in brain microglia (Butovsky et al., 2014; Mildner et al., 2017), and in those expressing the pan-neuronal marker *RBFOX3*. We selected the ACC, the dlPFC, the ventromedial prefrontal cortex (vmPFC), the caudate nucleus, the dentate gyrus (DG) of the anterior hippocampus, and the cerebellum (crus I lobule) as regions of interest for single-molecule FISH experiments (Supplementary Table 1).

We found *BMAL1* expression to strongly map to *RBFOX3*-enriched areas, such as the cortical layers, as well as the granule cell layer of the DG and cornu ammonis (CA) regions of the hippocampus (Supplementary Figure 1), suggesting that the core clock gene is preferentially expressed in grey matter regions. Following quantification of *BMAL1*^+^ cells in the various brain regions (Figure 1A-B), we found that the grey matter of the ACC exhibited on average the highest percentage of cells expressing *BMAL1* (21.49±1.72 %), whereas the dentate gyrus (5.67±0.85%) and the cerebellar cortex (5.64±1.15 %) had the lowest percentages (Figure 1B; Supplementary Table 2). A significant correlation was found between the percentage of cells expressing *BMAL1* and the post-mortem interval between death and refrigeration of the body (refrigeration delay) (Supplementary Figure 2A-D). We also found a significant interaction between the proportion of *BMAL1*^+^ cells and time of death (Supplementary Figure 2E).

**Figure 1.**
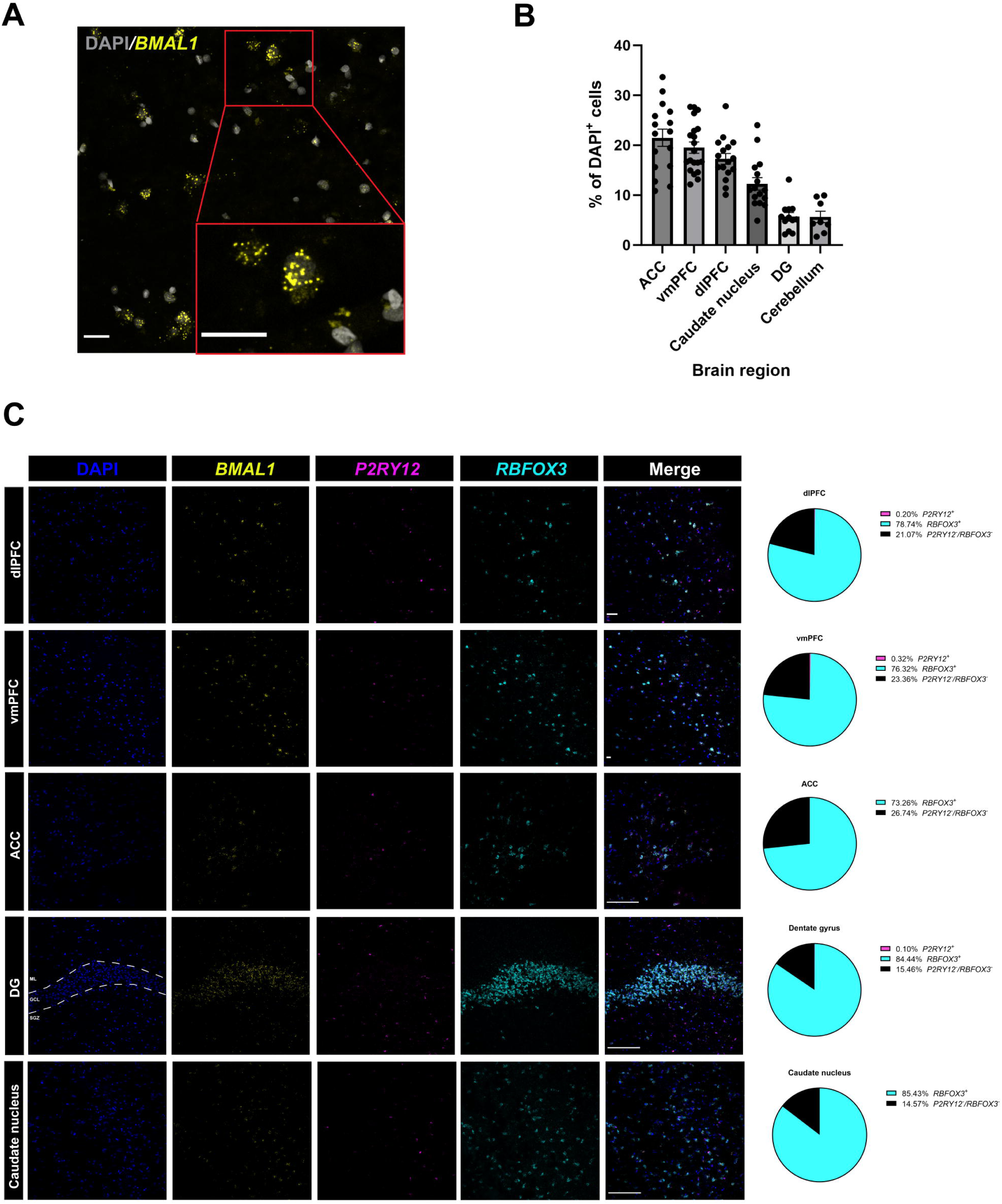
*BMAL1*^+^ cells mostly express *RBFOX3*, while very few express *P2RY12* in the adult human brain. (A) Representative confocal image of *BMAL1*^+^ cells in the dlPFC. Scale bars = 50 µm (scale bar of expanded inset = 50 µm). (B) Percentage of DAPI^+^ cells expressing *BMAL1* in the dlPFC grey matter (n=4), vmPFC grey matter (n=5), ACC grey matter (n=4), DG (n=3), caudate nucleus (n=4), and cerebellum (crus I) (n=2) of non psychiatric sudden-death individuals. Each dot corresponds to one region of interest (ROI) (from one staining replicate per subject). Bar plot shows mean with SEM. (C) Representative confocal images of *BMAL1*^+^ cells expressing *RBFOX3* and *P2RY12* microglia in the different brain regions. The corresponding pie charts demonstrate the average percentage of *BMAL1*^+^ cells expressing *RBFOX3* or *P2RY12* in the dlPFC grey matter (n=4), vmPFC grey matter (n=5), ACC grey matter (n=4), DG (n=3) and caudate nucleus (n=4). For each subject, four ROIs were counted. Scale bars = 50 µm. Dotted lines delineate the GCL from the underlying SGZ and the ML. dlPFC = dorsolateral prefrontal cortex; vmPFC = ventromedial prefrontal cortex; ACC = anterior cingulate cortex; DG = dentate gyrus of the hippocampus; GCL = granule cell layer, ML = molecular layer, SGZ = subgranular zone. Source data is found in Supplementary Table 2.

When comparing the percentage of *BMAL1*^+^ cells expressing *RBFOX3* or *P2RY12*, strikingly, we found that the majority of *BMAL1*^+^ cells expressed *RBFOX3,* which was consistent in all brain regions (average of 79.35±1.51%) (Figure 1C), but the cerebellum (Figure 2; Supplementary Figure 2F; Supplementary Table 2), thus revealing that the majority of *BMAL1*-expressing cells represent neurons in the human brain. In contrast, on average, less than 1% (0.15± 0.09%) of *BMAL1*^+^ cells expressed *P2RY12* in all brain regions (Supplementary Figure 2F) and most ROIs quantified contained extremely low percentages of *P2RY12*^+^ microglia expressing *BMAL1* (Supplementary Figure 2G), most of them reaching zero, thus indicating that only a very small subset of microglia, if any, expresses the core clock gene.

**Figure 2.**
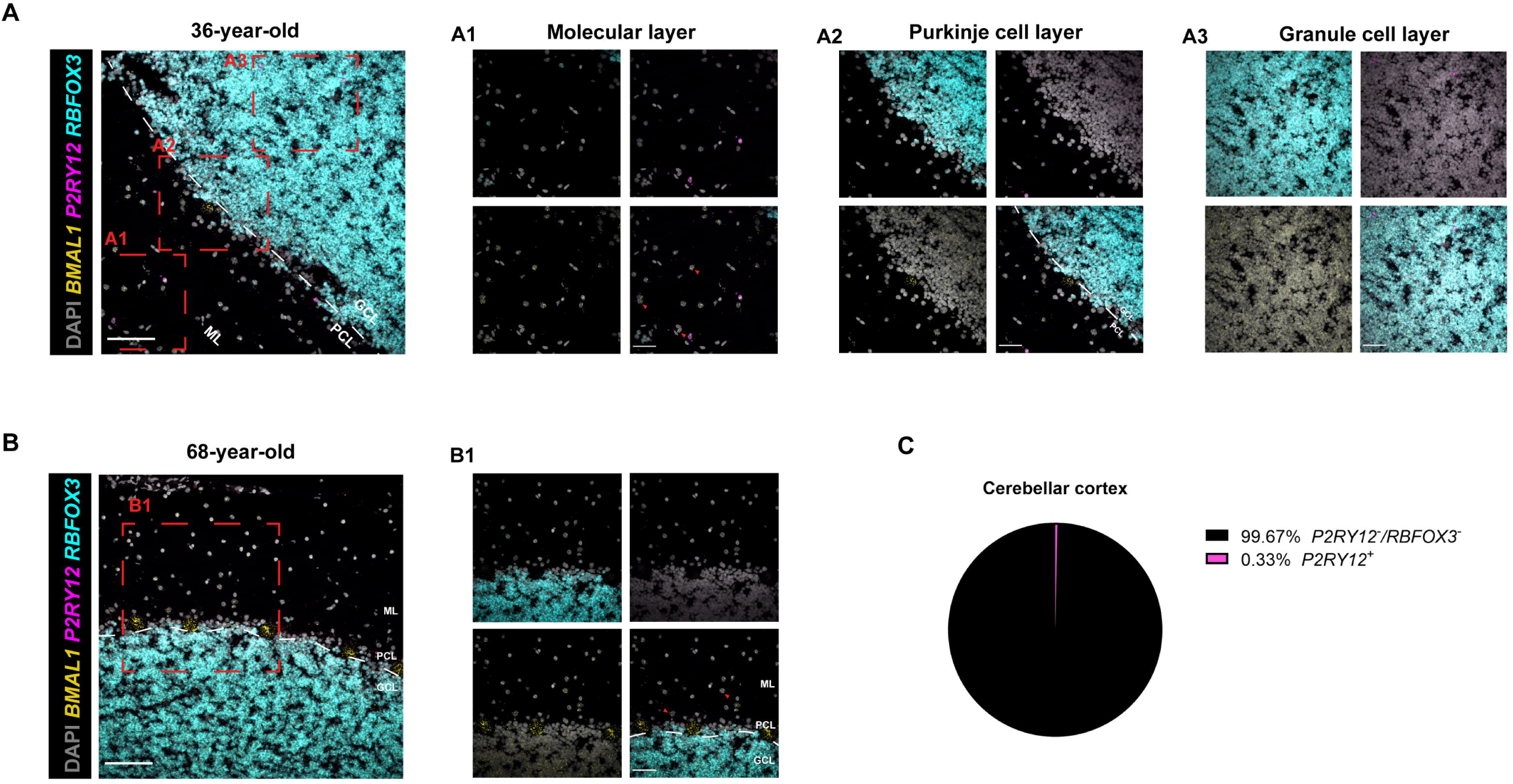
*BMAL1*^+^ cells are mostly devoid of *RBFOX3* expression and very few express *P2RY12* in the adult human cerebellum. (A) Representative confocal image of *BMAL1* expression in the cerebellar cortex (crus 1 lobule) with higher magnification images in the ML (A1), PCL (A2), and GCL (A3) of a 36-year-old sudden-death individual. Scale bars = 50 µm. (B) Representative confocal image of *BMAL1* expression in the cerebellar cortex (crus 1 lobule) with higher magnification images of all three layers in a 68-year-old sudden-death individual (B1). Scale bars = 50 µm. Red arrows point to *BMAL1*^+^*RBFOX3*^−^ cells in the molecular layer. (C) Pie chart demonstrating the average percentage of *BMAL1*^+^ cells expressing *RBFOX3* or *P2RY12* in the cerebellum. For each subject (n=2), four ROIs were counted. Dotted lines delineate the PCL from the GCL and ML. GCL, granule cell layer; PCL, Purkinje cell layer; ML, molecular layer. Source data is found in Supplementary Table 2.

In the human cerebellum (crus I lobule), strikingly, we found a substantial number of *BMAL1*^+^ cells without expression of *RBFOX3* or *P2RY12* (average of 99.67±0.33 %) (Figure 2A-C; Supplementary Table 2), almost all of them originating from the molecular layer of the cerebellar cortex (Figure 2A-B). We speculate that these cells may represent stellate cells or basket cells, as they are known to not express RBFOX3 (Weyer & Schilling, 2003). We also noted many *BMAL1* transcripts in *RBFOX3*^−^ Purkinje cells, whereas the granule cell layer mostly contained *BMAL1* mRNA puncta widely distributed throughout it (Figure 2A-B). This finding corroborates previous observations of higher number of *Bmal1* transcripts in Purkinje cells compared to granular neurons of the cerebellum in a single-cell RNA sequencing study of the mouse brain (Saunders et al., 2018; Zheng et al., 2023).

No significant correlation was found between the percentage of *BMAL1*^+^ cells expressing *RBFOX3* and the different covariates (Supplementary Figure 3A-D). In addition, we did not find a significant interaction between the proportion of *BMAL1*^+^*RBFOX3*^+^ cells and time of death (Supplementary Figure 3E). Similarly, the percentage of *BMAL1*^+^ cells expressing *P2RY12* did not correlate with any of the covariates (Supplementary Figure 4A-D) and was not associated with time of death (Supplementary Figure 4E).

We next characterized BMAL1 protein expression in microglia by isolating them from whole mouse brain (the olfactory bulb and cerebellum were excluded) samples and from fresh post-mortem human vmPFC grey matter with CD11b magnetic beads using Magnetic-Activated Cell Sorting (MACS). It was previously shown that CD11b positive selection using MACS columns produces 90% enrichment of CD11b^+^ microglia (Rayaprolu et al., 2020). More recently, in human post-mortem vmPFC samples, we showed that the majority of CD11b positive cells are resident microglia expressing P2RY12 (Belliveau et al., 2024). Furthermore, CD45 high brain infiltrating macrophages represent less than 5% of cells within CD11b^+^ brain myeloid cells and therefore, MACS reliably enriches for CD11b^+^ brain myeloid cells (Rangaraju et al., 2018). Our LC-MS proteomics analysis of CD11b^+^ isolated cells showed a lack of expression of core clock components (BMAL1, CLOCK, PERs, and CRYs) in microglia (Supplementary Table 3). Considering that these circadian clock markers correspond to cytoplasmic or nuclear but not membrane-bound proteins, they are expected to be present among the detected peptides. The list of detected proteins and their abundance in mice and humans can be found in Supplementary Table 3.

We further explored the possibility of other cell types expressing *BMAL1* by focusing on the dlPFC. Using an RNAscope probe directed against the astrocytic marker *ALDH1L1* and the oligodendrocyte precursor cell (OPC) marker, *PDGFRA*, in dlPFC sections, we qualitatively assessed *BMAL1* expression in these cell types. We noted *BMAL1* expression in a few *ALDH1L1*^+^ cells but could not easily identify *BMAL1^+^PDGFRA^+^* cells, possibly indicating higher *BMAL1* expression in astrocytes compared to OPCs, at least in the human dlPFC (Figure 3A). When quantifying *ALDH1L1* and *RBFOX3* expression on the same dlPFC sections, on average, 2.95±0.87% of *BMAL1*+ cells expressed *ALDH1L1*, while the majority expressed *RBFOX3* (59.28±3.88%) (Figure 3B; Supplementary Table 4), further revealing that a small subset of *BMAL1* expressing cells is astrocytic. In addition, on average, we found 15.87 ± 4.79% of *ALDHL1L*^+^ cells expressing *BMAL1,* which was significantly lower than the number of astrocytes negative for *BMAL1* (Figure 3C; Supplementary Table 4). We would like to note that while the majority of *BMAL1*^+^ cells are neuronal, the remaining percentage of cells that do not express either *RBFOX3*, *ALDH1L1*, *PDGFRA* and *P2RY12* may represent other cells, such as endothelial cells and mature OPCs, and other RBFOX3^−^ neuronal cell types, which warrants further investigation.

**Figure 3.**
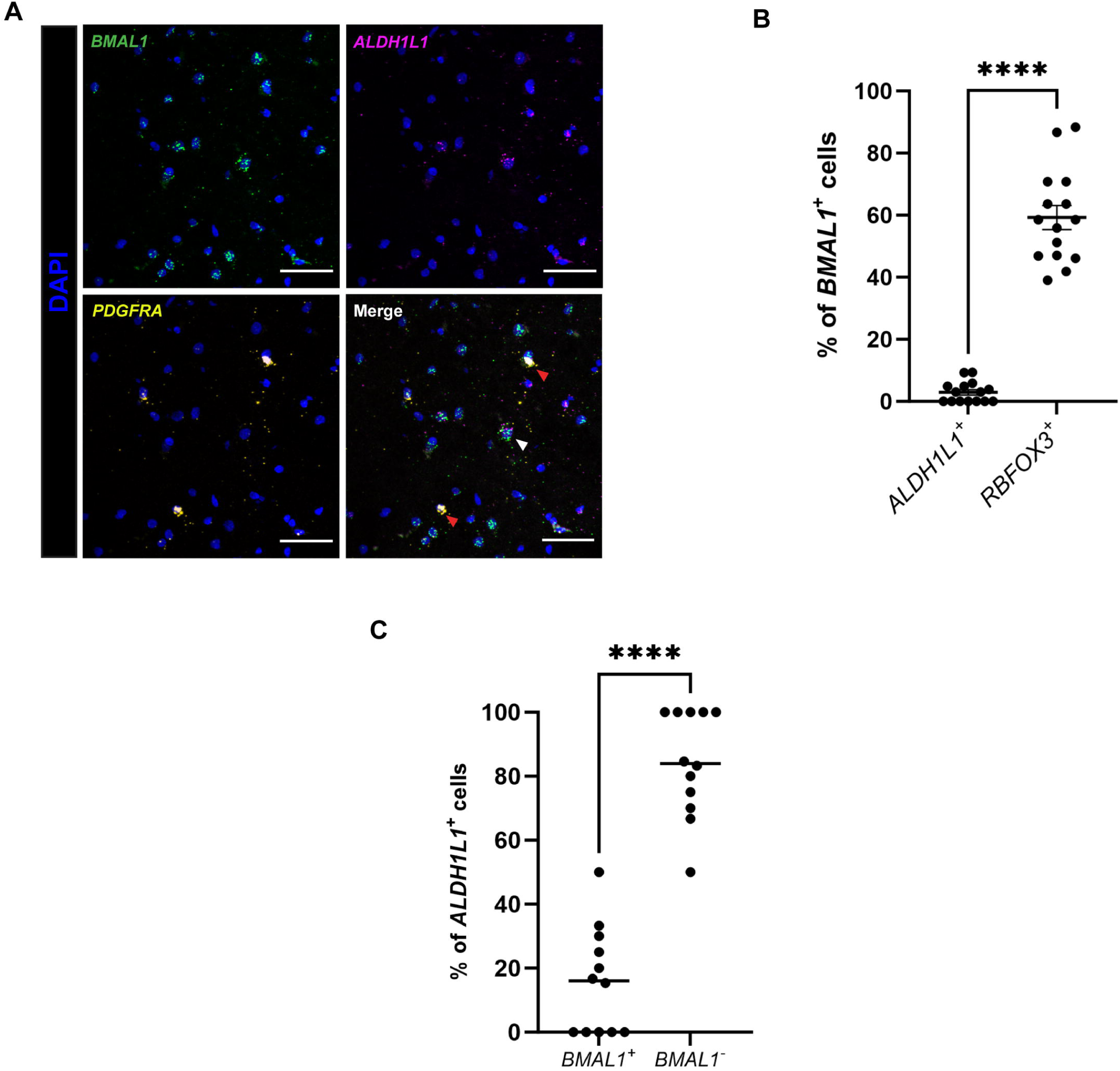
*BMAL1* is expressed in *ALDH1L1*^+^ cells in the adult human dlPFC. (A) Representative confocal images of cells co-expressing *BMAL1* and *ALDH1L1* in the dlPFC of a sudden-death individual. White arrows point to *BMAL1*^+^*ALDH1L1*^+^ cells. Red arrows point to *PDGFRA*^+^ cells without *BMAL1* expression. Scale bars = 50 µm. (B) Percentage of *BMAL1*^+^ cells expressing *ALDH1L1* or *RBFOX3* in the dlPFC (two-tailed Student t-test, p < 0.0001) (3 subjects; 4 ROIs/subject). Each dot corresponds to one region of interest (ROI) (from one staining replicate per subject). Scatter plot shows mean with SEM. (C) Percentage of *ALDH1L1*^+^ cells expressing or not *BMAL1* in the dlPFC (two-tailed Student t-test, p < 0.0001) (3 subjects; 4 ROIs/subject). Each dot corresponds to one region of interest (ROI) (from one staining replicate per subject). Bar plot shows mean with SEM. **** p< 0.0001 in two-tailed Student t-test. Source data is found in Supplementary Table 4.

We next questioned whether our RNAscope and proteomic findings in the human brain could be replicated using tissue from mouse brain. To this end, we immunostained brain sections using an antibody against Bmal1 which has been validated in Bmal1^−/−^ mice demonstrating excellent specificity (Chu et al., 2013). Remarkably, Iba1 positive cells appeared completely devoid of Bmal1 signal (green) in cortex, corpus callosum, and striatum (Supplementary Fig. 5A,B,D), while GFAP-expressing cells co-stained for Bmal1 (Supplementary Fig. 5C). These findings strongly argue that microglia at least in the mouse cortex and striatum do not express the essential clock component Bmal1, while astrocytes do, consistent with their role as circadian rhythm generators (Brancaccio et al., 2019).

### Analysis of single-nucleus RNA-sequencing and spatial transcriptomic datasets reveals BMAL1 enrichment in neuronal populations in the human hippocampus and dorsolateral prefrontal cortex of non-psychiatric sudden-death individuals

Findings from a recent analysis of a single cell RNA-sequencing (scRNA-sequencing) study of the mouse brain (Saunders et al., 2018) suggest that *BMAL1* expression is more highly enriched in neuronal populations, including both excitatory and inhibitory neurons, than in microglia in various brain regions, including the hippocampus and the frontal cortex (Zheng et al., 2023). To further explore differences in *BMAL1* expression between neuronal cell types and microglia in the rodent and human brain, we utilized publicly available snRNA-sequencing datasets of the mouse prefrontal cortex (PFC) and hippocampus (Habib et al., 2017), the human hippocampus (Ayhan et al., 2021), and the human dlPFC (Maitra et al., 2023). In addition to *BMAL1* (*ARNTL*), we also assessed the expression of other core clock genes, more specifically *DBP*, *CRY1*, *PER1* and *PER2*. Similar to our RNAscope findings, we found neuronal clusters (excitatory and/or inhibitory) in all datasets to prominently feature *Bmal1/BMAL1* transcripts compared to microglial clusters, which had only 1-7% of nuclei with non-zero expression levels of *BMAL1* (Figure 4A-C; Supplementary Table 5). For instance, in Habib et al. (2017), around 19% of nuclei in the exPFC8 cluster, corresponding to excitatory prefrontal cortical neurons, expressed higher levels of *BMAL1*. This was also true for about 17% of the nuclei of the inhibitory neuron-presenting In2 cluster of Ayhan et al. dataset and 30% of the nuclei from the excitatory neuronal cluster reported in Maitra et al. (2023).

**Figure 4.**
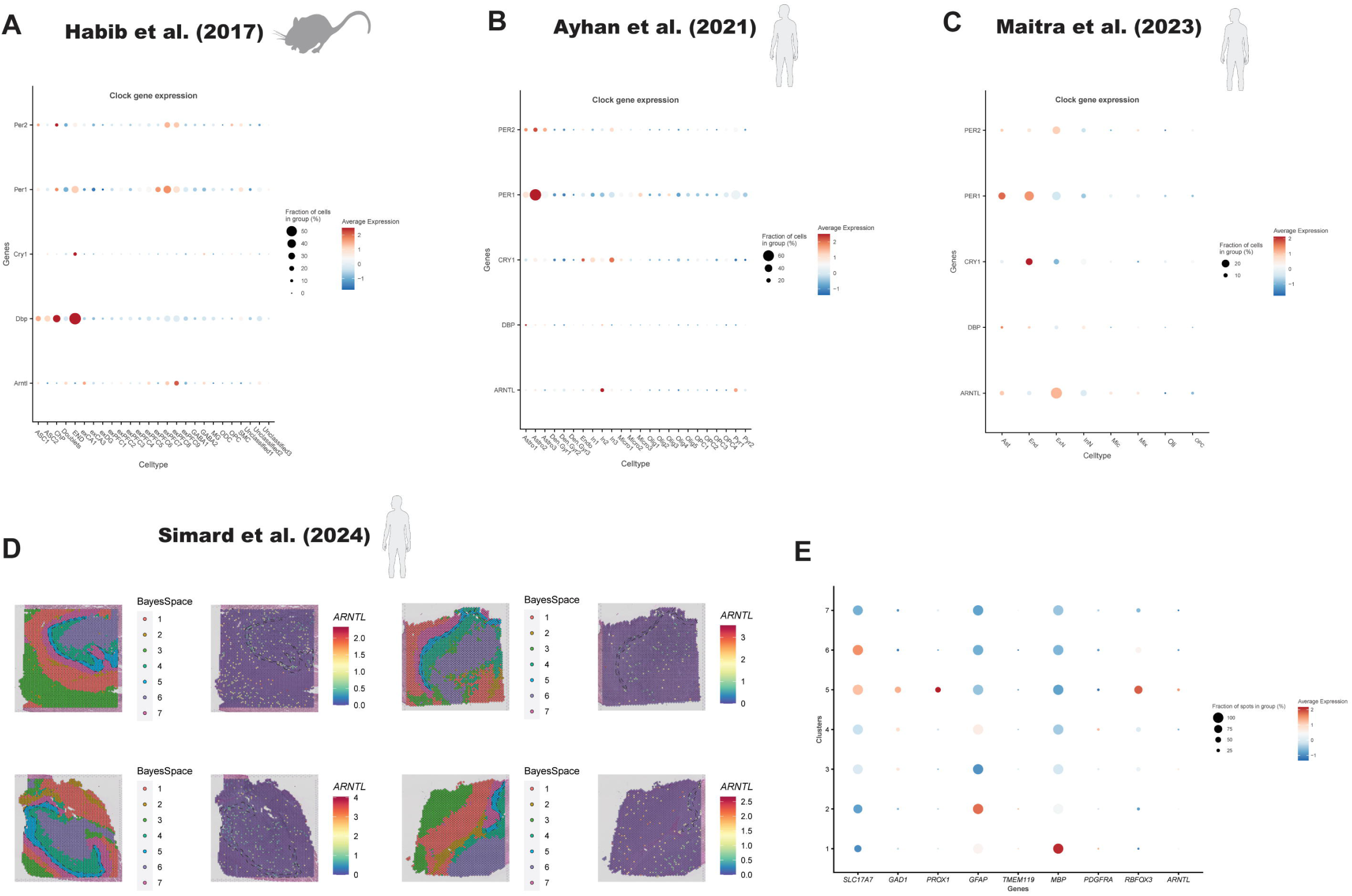
Circadian clock gene expression in external single-nucleus RNA sequencing and spatial transcriptomic datasets. Dot plot showing scaled average expression levels of *ARNTL (BMAL1)*, *DBP*, *CRY1*, *PER1,* and *PER2* in cell type specific clusters in a single-nucleus RNA sequencing dataset of the mouse PFC and hippocampus (Habib et al., 2017) (A), the hippocampus (Ayhan et al., 2021) (B), and the human dlPFC (Maitra et al., 2023) (C). Red (blue) hues indicate higher (lower) expression levels. Source data is presented in Supplementary Table 5. (D) Spatial plots showing BayesSpace clustering and log-normalized expression of *ARNTL* in the hippocampus of 4 non-psychiatric sudden-death individuals (4 males) (Simard et al., 2024). Dotted lines were added to delineate the granule cell layer of the DG. Clusters are annotated as follows in Simard et al. (2024): Cluster 1 = white matter; Cluster 2 = blood vessels; Cluster 3 = CA1 and subiculum; Cluster 4 = polyform cell layer and subgranular zone; Cluster 5 = granule cell layer; Cluster 6 = CA4 and CA3; Cluster 7 = molecular layer. (e) Dotplot showing scaled average expression levels of canonical cell type markers (*SLC17A7, GAD1*, *PROX1*, *GFAP*, *TMEM119*, *MBP*, *PDGFRA,* and *RBFOX3*) and *ARNTL* in each BayesSpace cluster. Around 16% of spots in Cluster 5 (granule cell layer), enriched for neuronal markers, show higher *ARNTL* expression levels compared to other clusters. Source data is presented in Supplementary Table 6.

Furthermore, we also found lower expression levels of *DBP*, *CRY1*, *PER1* and *PER2* in microglia clusters compared to other non-neuronal clusters, comprising astrocytes, and endothelial cells, in all three datasets, underscoring a microglia-specific low abundance of clock gene transcripts (Supplementary Table 5). Notably, we also found a very low account of *BMAL1* in OPC and oligodendrocyte clusters in all three datasets, with approximately only 1-4% of nuclei expressing low levels of *BMAL1*, corroborating our RNAscope findings.

We next sought to use a spatial transcriptomic dataset of the human hippocampus, previously generated by our group (Simard et al., 2024). We found that approximately 16% of spots in Cluster 5, the *PROX1*-expressing granule cell layer, were more highly expressing *BMAL1* compared to other *RBFOX3*-enriched clusters, such as Clusters 3, 4, and 6 (Figure 4D-E, Supplementary Table 6). In agreement with our analysis of single-nucleus RNA sequencing datasets, we found lower *BMAL1* expression in the *MBP*-expressing Cluster 1 in the hippocampal sections, further revealing low expression of the core clock gene in white matter regions of the brain. While previous research has shown that OPC proliferation and oligodendrocyte maturation may be dependent on Bmal1 expression (Dean et al., 2022) and the entire circadian clock machinery (Colwell & Ghiani, 2020), more recent evidence in mice reveals that the involvement of Bmal1 in OPC maintenance may be less prominent in adulthood, but crucial during early developmental oligodendrogenesis (Rojo et al., 2023). However, as we did not quantify the number of *BMAL1*^+^*PDGFRA*^+^ cells in our FISH experiments, we cannot exclude the possibility of a small population of OPCs expressing the core clock gene.

### Analysis of a single-nucleus ATAC sequencing dataset reveals lower BMAL1 chromatin accessibility in microglia compared to neuronal populations in non-psychiatric sudden-death individuals

To decipher potential regulatory mechanisms that may explain the unexpectedly lower gene expression of *BMAL1* in microglia, we utilized snATAC-sequencing chromatin accessibility data from the postmortem dlPFC of healthy individuals (n=40) (Chawla et al., 2023). Our results show that chromatin accessibility measured at the *BMAL1* gene promoter (2kbp upstream of the transcription start site (TSS)) showed significantly lower access in microglia compared to neurons One-way ANOVA, Tukey HSD p<0.0001) (Figure 5A). A similar pattern was observed for chromatin accessibility across upstream *cis*-regulatory regions of *BMAL1* (5kbp upstream of TSS with an exponential decay model) (Corces et al., 2020) (Figure 5B). These results suggest a strong role of upstream epigenetic factors in driving an overall lower expression of *BMAL1* in microglia compared to other cell classes, except oligodendrocytes. To further explore this pattern, we asked whether any open chromatin regions (OCRs) or cell-type-enriched *cis*-regulatory regions (marker cCREs, highest accessibility in one versus all other cell-types) were present in the *BMAL1* gene locus (50kbp upstream & downstream of TSS). Both microglial specific OCRs and marker cCREs were sparingly present in this locus compared to other cell classes except oligodendrocytes (Figure 5C-D, orange label). Furthermore, examining potential gene-regulatory linkages (snATAC-sequencing & snRNA-sequencing based peak accessibility and gene expression correlations, Pearson’s r>0.5) revealed many distal regulatory regions (e.g. enhancers) driving differential expression of *BMAL1* in neurons compared to microglia (Figure 5C-D). Together, these results suggest a role of cell-type specific epigenetic mechanisms in driving lower expression of *BMAL1* in microglia.

**Figure 5.**
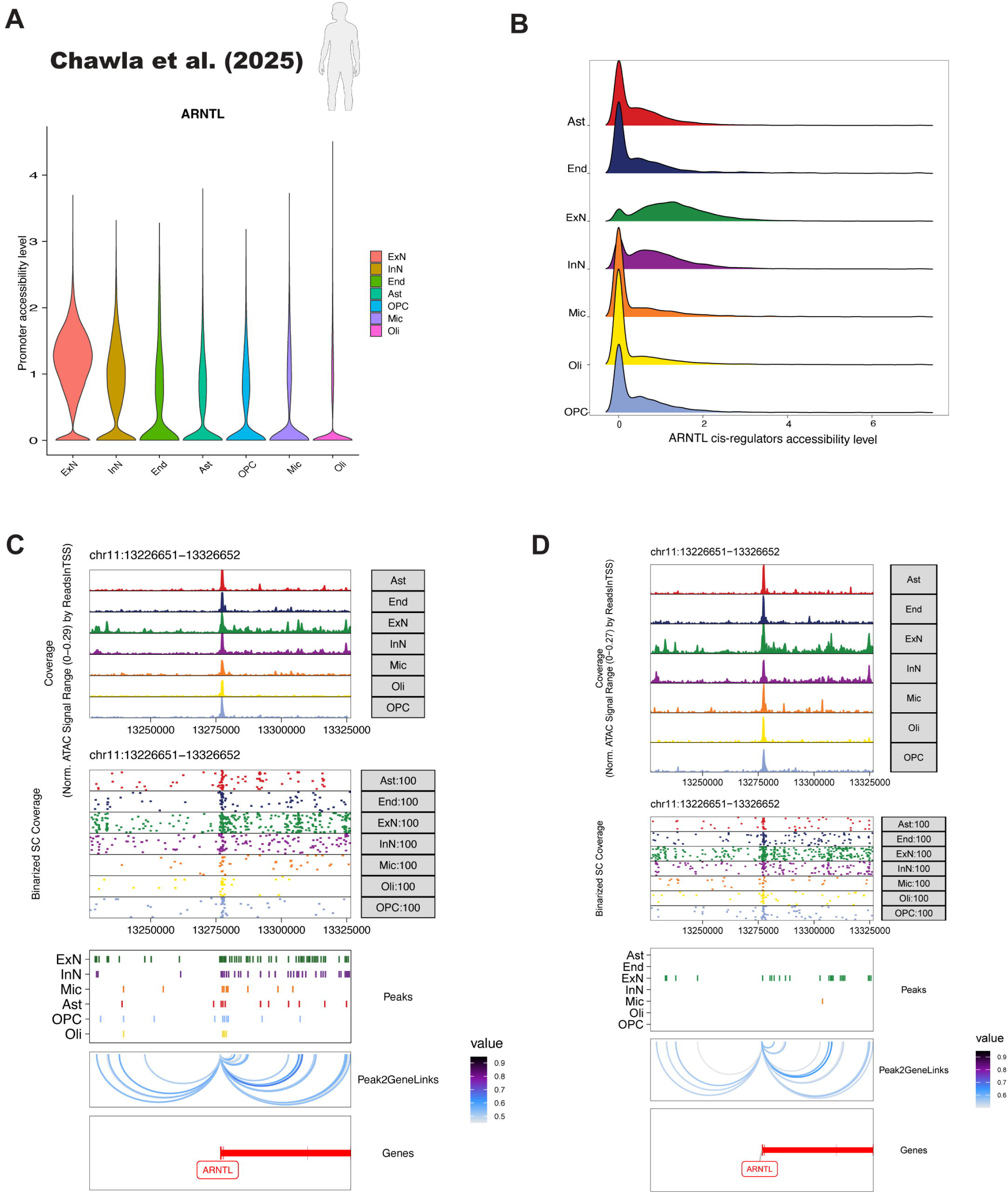
ARNTL gene accessibility in external single-nucleus ATAC sequencing dataset. (A) Violin plot shows promoter (2kbp upstream of TSS) accessibility for ARNTL (BMAL1) gene, ordered by average expression of ARNTL in each cell type from Chawla et al. (2025). One-way ANOVA with Tukey HSD was performed for pairwise comparison between cell types. (B) Ridge plots show chromatin accessibility across cis-regulatory regions of ARNTL (5kbp upstream of TSS using an exponential decay model) (Corces et al., 2020). (C-D) Multi-modal plots shows chromatin accessibility in each cell type (normalized by reads in TSS) at ARNTL gene locus (50kbp upstream and downstream of ARNTL TSS), followed by binarized single-cell coverage for 100 sampled cells from each cell type, and (C) OCRs in each cell type in this locus and (D) marker cCREs for each cell type in this locus. Finally, we show peak-to-gene linkages (Pearson’s r>0.5) between *ARNTL* gene-expression and their regulatory peaks in this locus.

Finally, in order to identify TFs that may be regulating *BMAL1* expression in microglia, we focused on microglial OCRs (up to +/− 500kbp of *BMAL1* TSS), whose chromatin accessibility positively correlated with *BMAL1* gene expression (peak-to-gene links, Pearson’s correlation, FDR<0.05). These microglial OCRs were found to be mainly present in intronic regions of *BMAL1* and promoter or distal regions of *LINC00958* but were absent from promoter regions of *BMAL1* (Supplementary Table 7). Most TF binding sites overrepresented in these regions (Homer, BH-corrected p<0.05) were ETS domain factors (such as ELF5, SPIB), which often bind to the E-box motifs in DNA (Supplementary Figure 6A).

Although *BMAL1* is lowly expressed in microglia, its open chromatin regions were not completely devoid of BMAL1 TF binding sites (Supplementary Figure 6B). This may suggest that, whereas the overall expression of *BMAL1* is low in microglia, its reduced expression may still have some downstream gene-regulatory effects due to the presence of BMAL1 TF binding sites in the open chromatin of microglia. To understand the downstream gene-regulatory effects of BMAL1 TF binding in microglia, we retained those BMAL1 binding sites which overlapped microglial OCRs and were linked to most likely effected genes (peak-to-gene correlations, Pearson’s r>0.5). As expected, our results showed an enrichment for genes regulating cell-activation and immune-related functions in microglia (Supplementary Figure 6C; Supplementary Table 8), in contrast to genes involved in neuron projection and synaptic activity, which were found enriched for excitatory neuronal OCRs (Supplementary Figure 7).

We also investigated the expression of *Bmal1* in mouse microglia using microarray findings with regard to the microglial translatome in Rahimian et al. (2024) and Boutej et al. (2017). Using an *in vivo* model system for analysis of the dynamic translational state of microglial ribosomes, with mRNAs as input and newly synthesized peptides as an output, both studies reported the dysregulation of several immune genes following brain ischemia and LPS administration (Boutej et al., 2017; Rahimian et al., 2024). Intriguingly, their data showed expression of *BMAL2,* the paralog of *BMAL1* which can substitute for Bmal1 in driving circadian rhythms (Shi et al., 2010), but revealed a lack of Bmal1 mRNA in microglial translating ribosomes. Based on these results and our own post-mortem human and mouse findings, we next explored the spatial expression of *BMAL2* in human post-mortem dlPFC to determine whether its expression may compensate for low *BMAL1* expression in microglia.

Notably, similar to BMAL1, we observed a significantly lower accessibility for BMAL2 promoter and cis-regulatory regions in microglia compared to neurons (Supplementary Figure 8A-E). We next explored how *BMAL2* is spatially expressed in the dlPFC (n=3) by using RNAscope, and following quantification, we found that, on average, more than half (53.00±16.39%) of *BMAL1*^+^*P2RY12*^−^ cells expressed *BMAL2* in the dlPFC of non-psychiatric individuals (Supplementary Figure 8F-G). In contrast, *BMAL1* and *BMAL2* co-expression was seemingly lacking in the *P2RY12*^+^ resident microglia. Considering the high percentage of *BMAL1*^+^ cells expressing *RBFOX3*, these observations further reveal that *BMAL1* and *BMAL2* co-expression is mainly present in neuronal cells. Overall, our findings do not support a role of BMAL2 as a functional substitute for BMAL1 in microglia.

## Discussion

In the current study, we used a variety of methods to gain insights in the expression pattern of the intrinsic clock gene *BMAL1* in the brain at the proteomic, transcriptomic, and epigenomic levels. In summary, the data presented here indicates a strong enrichment for *BMAL1* expression in grey matter regions and very minimal expression of BMAL1 in microglia in the human brain, especially when compared to other neuronal cell populations. Moreover, mouse microglia, at least in cortex and striatum, seem completely devoid of Bmal1. Interestingly, a recent report revealed a significant decrease in Bmal1 expression in aged microglia of 21- to 23-month-old mice, corresponding to late adulthood in humans, compared to 6-7 month-old mice, representing middle adulthood in humans, suggesting a possible role of physiological aging for microglial suppression of Bmal1 (Sheehan et al., 2024). However, in our IF experiments, we observed that in 8–12-week-old mice, Bmal1 expression was virtually absent in microglia, arguing against intrinsic circadian rhythm generation in microglia even at a very young age. We also did not find correlations between BMAL1 gene expression or accessibility and age of death of individuals in the human data set of Chawla and colleagues’ (2023) (Supplementary Figure 9). In addition, using FISH on samples from adults between the ages of 25 and 78 years, we could demonstrate minimal *BMAL1* expression in microglia which did not correlate with age. In further support of our findings, a recent scRNA-sequencing study on human dIPFC showed lower *BMAL1* expression in microglia of the dlPFC during young, middle and late adulthood, compared to neuronal populations (Yang et al., 2024).

An overall low expression of BMAL1 in microglia can be explained by a lack of accessibility or openness of BMAL1 promoter or distal regulatory regions in microglia compared to neurons. We also identified several ETS domain TFs that may play a role in suppressing constitutive expression of BMAL1 in microglia. ETS domain TFs have been previously associated with regulating circadian transcription (Kim & Lazar, 2020) and are crucial for microglial functions (Yeh & Ikezu, 2019). It is also possible that constitutively *BMAL1* expression is suppressed in microglia by means of the identified TFs and other regulatory mechanisms must take place for allowing a rhythmic expression.

Our findings challenge the prevailing view that microglia, much like astrocytes and neurons, generally possess an intrinsic circadian clock. Different from most studies that assessed clock gene expression in microglia (Fonken et al., 2015; Hayashi et al., 2013; Nakazato et al., 2017; Nakazato et al., 2011; Ni et al., 2019; Wang et al., 2021), we queried frozen or fixed intact tissue using *in situ* hybridization or immunohistochemistry. Tissue intactness and immediacy of expression analysis post tissue retrieval might be critical to preserve the native physiological state, which could possibly alter when microglia first undergo an isolation process from intact tissue prior to assessment. Furthermore, isolated cell populations may suffer from cellular impurities and this might be a particular concern for microglia as they phagocytose other cells (Kamei & Okabe, 2023), which may not be dissociable when isolating microglia from brain tissue. Such impurities could potentially give rise to detectable levels of Bmal1 and other clock components, which might even be rhythmic, when employing very sensitive expression analysis techniques such as quantitative RT-PCR, a method commonly utilized to demonstrate rhythmic clock gene expression in microglia (isolates) (Fonken et al., 2015; Nakazato et al., 2011; Wang et al., 2020). In further support of a lack of clock gene expression, and thus an intrinsic rhythm generating capacity in microglia, we failed to detect Bmal1 in mouse microglia employing an Bmal1 antibody whose specificity has been validated in *Bmal1^−/−^* mice (Chu et al., 2013).

What could be the functional significance of low or absent *BMAL1* expression in microglia? Clearly, considering that Bmal1 is an absolutely essential clock component (Bunger et al., 2000), our data would argue that microglia by and large don’t harbor a functional circadian clock, which is in contrast to most other cell types across the body axis (Mannic et al., 2013; March et al., 2024; Pulimeno et al., 2013). From an evolutionary perspective, a time-of-day difference in cellular physiology, which is the consequence of a ticking intrinsic circadian clock, could prevent microglia to optimally respond to brain insults that are extrinsic or intrinsically born such as pathogen infection or stroke, which may strike at anytime during the 24-hour solar day cycle. In case of such insults which may lead to brain ischemia and apoptotic cell death, microglia multiply rapidly and secrete a variety of growth factors and cytokines to counteract stroke-related neuronal damage (Lalancette-Hébert et al., 2007). Specifically, the need to proliferate rapidly upon insult might have led to the selective suppression of clock function in microglia during evolution, given the evidence that clock components can act as tumor suppressors (Battaglin et al., 2021).

It is also worth noting that *BMAL1* epigenetic alterations contribute to the development of hematologic malignancies by disrupting the cellular circadian clock (Taniguchi et al., 2009). The *BMAL1* gene is transcriptionally silenced by promoter CpG island hypermethylation in hematologic malignancies such as diffuse large B-cell lymphoma, and acute lymphocytic and myeloid leukemias (Taniguchi et al., 2009). Indeed, *BMAL1* depletion caused by RNA interference in unmethylated cells promotes cell proliferation and tumor growth (Taniguchi et al., 2009). Therefore, it is possible that, in addition to the accessibility landscape, other epigenetic factors, such as methylation, may be responsible for reducing *BMAL1* expression in microglia.

The study was limited by small sample sizes for the RNAscope and proteomic experiments and the age range of the subjects. While we included subjects from young to late adulthood, we did not assess the phenotype of *BMAL1-*expressing cells in adolescents and infants, which should be investigated in future research. In addition, while we looked at the relationship between *BMAL1* expression in microglia and time of death in our post-mortem human samples, we did not explicitly explore *BMAL1* expression at different time points using an animal model and therefore, could not provide information on how BMAL1 expression fluctuates with time. It should be however noted that Bmal1 expression levels are not known to drop to non-detectable levels at any given time of day in any brain regions examined. Thus, the total absence of Bmal1 immunoreaction in mouse microglia seems incompatible with normal Bmal1 protein fluctuation and instead appears more consistent with a general absence of Bmal1 from microglia in the intact tissue context. Future studies should also compare *BMAL1* expression in different neuronal subtypes (e.g. excitatory and inhibitory), as well as assess differences in expression in the context of neurodegeneration and neuropsychiatric conditions, such as major depressive disorder and schizophrenia, disorders where Bmal1 function may be implicated (Valeri et al., 2022; Zheng et al., 2023). We also did not investigate possible sex differences in the expression of *BMAL1* in neuronal and microglial cell types as our sample sizes did not allow us to conduct these analyses. Notwithstanding these limitations, our study provides important and novel insight into the spatial distribution of *BMAL1* in various regions of the human brain. The current findings may prompt a fresh look at microglia versus neuronal physiology in the context of the daily light-dark cycle.

## Methods

### 1. Human post-mortem brain samples

Brain samples used in this study were obtained from the Douglas-Bell Canada Brain Bank (Montreal, Canada). In collaboration with the Quebec Coroner’s Office and with informed consent from next of kin, phenotypic information was obtained with standardized psychological autopsies. Presence of any or suspected neurological/neurodegenerative disorder signaled in clinical files constituted an exclusion criterion. Controls, which correspond to individuals who died by natural or accidental causes of death, without any brain disorder, are defined with the support of medical charts and coroner records. Toxicological assessments and medication prescription were also obtained. Detailed information on the samples used for multiplexed fluorescence *in situ* hybridization (RNAscope) experiments and microglia isolation pull-down is reported in Supplementary Table 1. Subjects for each brain region were matched for age, post-mortem interval (PMI) and refrigeration delay (delay between time of death and storage of the body at 4°C). Only pH values differed between brain regions.

### 2. Multiplexed fluorescence in situ hybridization (RNAscope) and data quantification

Human post-mortem frozen unfixed samples from the dlPFC, vmPFC, ACC, cerebellum (crus I lobule), caudate nucleus, and frozen fixed anterior hippocampal samples were cryosectioned at 10 μm, collected on Superfrost charged slides, and stored at −80°C. *In situ* hybridization was performed using Advanced Cell Diagnostics RNAscope® probes and reagents (ACD Bio, RNAscope® Multiplex Fluorescent v2 Assay) following the manufacturer’s instructions. Briefly, tissue sections were fixed in 10% neutral buffered formalin for 15 minutes at 4°C. A series of dehydration steps was next performed with different concentrations of ethanol baths (50%, 70%, 95%, 100%) and the sections were next air dried for 5 minutes. A hydrogen peroxide treatment (10 minutes at room temperature) was then done on the tissue sections. The sections were after incubated for two hours in a temperature-controlled oven (HybEZ II, ACDbio) using different probe combinations: **(1)** Hs-ARNTL (catalog #470091), Hs-P2RY12-C2 (catalog #450391-C2), Hs-RBFOX3-C3 (catalog #415591-C3); **(2)** Hs-ARNTL, Hs-P2RY12-C2, Hs-ARNTL2-C3 (catalog #450201-C3); **(3)** Hs-ARNTL, Hs-ALDH1L1-C2 (catalog #438881-C2), Hs-PDGFRA-C3 (catalog #604481-C3); **(4)** Hs-ARNTL, Hs-ALDH1L1-C2, Hs-RBFOX3-C3. After amplification steps (AMP 1–3), probes were fluorescently labeled with Opal Dyes (Opal 520 for Channel 1, Opal 570 for Channel 2, Opal 690 for Channel 3; Perkin Elmer; 1:300 dilution for each). Autofluorescence from lipofuscin was quenched using the reagent TrueBlack (Biotium) for 40 seconds and sections were finally coverslipped with Vectashield mounting medium containing 4′,6-diamidino-2-phenylindole (DAPI) to label the nucleus (Vector Laboratories, H-1800). Slides were stored at 4 °C until imaging. For the hippocampal sections, two additional steps were performed: a 30-minute incubation step (at 60°C) before formalin fixation and an antigen retrieval step before a 30-minute protease digestion at 40°C (Protease III, undiluted).

The slides were imaged at 20X magnification using the Olympus VS120 virtual slide scanner and the scans were transferred to QuPath (v.0.3.0) (Bankhead et al., 2017) for further analysis. Regions of interest (ROIs) were defined as 500 µm x 500 µm tiles and were created for each scan. Four ROIs were then randomly selected for each scan. For the dlPFC, vmPFC and ACC, the grey matter was first delineated using the neuronal marker *RBFOX3* staining. For the hippocampal sections, we focused on the dentate gyrus (DG), as this region was present in all sections. Therefore, the subgranular zone (50 µm from the inner granule cell layer), the granule cell layer (GCL), traced according to DAPI staining, and the molecular layer (200 µm from the outer granule cell layer) were individually defined, and the annotations were then merged. The 500 µm x 500 µm tiles were created in the resulting merged DG annotation. In the cerebellar sections, each ROI included the granule cell layer, the Purkinje cell layer, and the molecular layer. Cells were counted within the ROIs using StarDist (Schmidt et al., 2018) (0.5 for detection threshold and 0.2 µm/pixel for pixel size) for cerebellar sections as clumped cells were highly present in the granule cell layer and using QuPath’s automated cell detection based on DAPI staining for all other sections. Single measurement classifiers were created for *BMAL1*, *RBFOX3,* and *BMAL2,* and counts were manually verified. Cells expressing *P2RY12* or *ALDH1L1* were manually counted. Only cells with more than 4 puncta were included in the analysis. Source data is presented in Supplementary Table 2.

### 3. Confocal microscopy

Representative images of *BMAL1, BMAL2, P2RY12, RBFOX3, ALDH1L1* and *PDGFRA* expression in the different brain regions were taken using the Olympus FV1200 laser scanning confocal microscope with 20x (NA: 0.75) and 40x objectives (NA: 0.95). The XY axis pixel number (1024×1024), Kalman averaging (2), and laser scanning speed (4µs/pixel) were modified to these settings to improve image resolution. Laser power and detection voltage parameters were adjusted between subjects for each set of experiments to improve image quality. We optimized all parameters to decrease signal to noise ratio and minimize autofluorescence from lipofuscin and cellular debris.

### 4. CD11b positive microglia isolation pull-down

Fresh post-mortem brain tissue samples used for microglia isolation pull-down were acquired from the DBCBB. Upon arrival at the DBCBB, samples underwent immediate processing for CD11b+ microglia isolation, according to the following procedure. Grey matter was first dissected from 1 cm^3^ blocks of fresh non-frozen vmPFC. To obtain murine microglia, brains excluding the olfactory bulb and cerebellum were dissected from two male C57BL/6 mice (3-month-old) after perfusion with ice-cold PBS and then immediately subjected to the CD11b selection procedure. Microglia were isolated using the Miltenyi Biotec CD11b MicroBeads (cat # 130-093-634, Gaithersburg, MD, USA), which were developed for the isolation or depletion of human and mouse cells based on their CD11b expression. In humans, myeloid cells, such as cerebral microglia, strongly express CD11b, whereas NK cells and some activated lymphocytes weakly express the marker. The CD11b^+^ microglia were isolated using a combination of enzymatic and mechanical dissociations following instructions from Miltenyi Biotec. The Neural Tissue Dissociation Kit (Papain, cat # 130-092-628, Miltenyi Biotec, Gaithersburg, MD, USA) and the gentleMACS™ Dissociator (cat # 130-093-235, Miltenyi Biotec, Gaithersburg, MD, USA) are effective in generating single-cell suspensions from neural tissues prior to subsequent applications, such as MACS® Cell Separation. Following the mechanical and enzymatic dissociation steps, red blood cells were eliminated using Red Blood Cell Lysis Solution (cat # 130-094-183, Miltenyi Biotec, Gaithersburg, MD, USA). These steps were followed by application of Debris Removal Solution, which is a ready-to-use density gradient reagent (cat # 130-109-398, Miltenyi Biotec, Gaithersburg, MD, USA). This reagent allows for a fast and effective removal of cell debris from viable cells after dissociation of various tissue types, while applying full acceleration and full brake during centrifugation. Prior to the positive selection of CD11b^+^ cells, Myelin Removal Beads were employed for the specific removal of myelin debris from single-cell suspensions (cat # 130-096-433, Miltenyi Biotec, Gaithersburg, MD, USA). Then, the CD11b^+^ cells were magnetically labeled with CD11b MicroBeads and the cell suspension was loaded onto a MACS® Column, which was placed in the magnetic field of a MACS Separator. The magnetically labeled CD11b^+^ cells were retained on the column, while the unlabeled cell fraction, which was depleted of CD11b^+^ cells, ran through the column. The magnetically retained CD11b^+^ cells were finally eluted following removal of the column from the magnetic field. The isolated cells were kept in −80°C freezer for downstream proteomic experiments. Finally, in a recent study, our group confirmed that the majority of CD11b^+^ cells isolated from vmPFC grey matter are indeed resident microglia (80-90%), but a small proportion of peripheral monocytes-macrophages was nevertheless present (Belliveau et al., 2024). Therefore, based on these findings, we cannot exclude the presence CD11b^+^ cells that are not microglia in the CD11b^+^ isolated cells analyzed in this study.

### 5. Liquid chromatography-mass spectrometry (LC-MS) proteomics

For each sample, proteins were loaded onto a single stacking gel band to remove lipids, detergents, and salts. The gel band was reduced with DTT, alkylated with iodoacetic acid, and digested with trypsin. Extracted peptides were re-solubilized in 0.1% aqueous formic acid and loaded onto a Thermo Acclaim Pepmap (Thermo, 75uM ID X 2cm C18 3uM beads) precolumn, and then onto an Acclaim Pepmap Easyspray (Thermo, 75uM X 15cm with 2uM C18 beads) analytical column separation using a Dionex Ultimate 3000 uHPLC at 230 nl/min with a gradient of 2-35% organic (0.1% formic acid in acetonitrile) over 3 hours. Peptides were analyzed using a Thermo Orbitrap Fusion mass spectrometer operating at 120,000 resolution (FWHM in MS1) with HCD sequencing (15,000 resolution) at top speed for all peptides with a charge of 2+ or greater. The raw data were converted into *.mgf format (Mascot generic format) for searching using the Mascot 2.6.2 search engine (Matrix Science) against mouse protein sequences (Uniprot 2023) and a database of common contaminant proteins. The database search results were loaded onto Scaffold Q+ Scaffold_5.0.1 (Proteome Sciences) for statistical treatment and data visualization.

### 6. Animals

8–12-week-old male C57BL/6J mice (Strain #: 000664, The Jackson Laboratory) were used for immunohistochemical analysis of Bmal1 expression. Animals were housed under a 12 hour:12 hour light:dark cycle at room temperature and experiments were performed in accordance with the Canadian Council on Animal Care guidelines and approved by the local McGill University Animal Care Committee.

### 7. Mouse immunohistochemistry

Brains were perfused and immunostained as previously described (Chu et al., 2013). Briefly, mice were removed from their housing cages at 2-5 hours after lights on of the daily light:dark cycle, deeply anesthetized with ketamine (100mg/kg)/xylazine (10 mg/kg), and perfused transcardially with PBS followed by 4% paraformaldehyde (PFA). Brains were postfixed in 4% PFA for 24 hours and then incubated in 30% sucrose in PBS overnight. Brains were embedded in optimal cutting temperature compound (OCT, Tissue-Tek; Cedarlane), frozen using dry ice, and cut at 30 microns using a cryostat (Leica). Brain sections were collected on SuperFrost Plus glass slides (VWR). For immunolabeling, sections were rinsed in PBS, incubated in blocking solution (3% goat serum in 0.1% Triton X-100 in PBS) for 1 hr, followed by incubation with primary antibodies in blocking solution at room temperature overnight. Sections were incubated in secondary antibodies for 2 hours at RT. Primary antibodies were used at the following dilutions: rabbit anti-Bmal1 (1:1000; cat# NB100-2288, Novus Biologicals), mouse anti-GFAP (1:500, EMD Millipore, cat# MAB360), mouse anti-Iba-1 (1:500, EMD Millipore, cat#MABN92). Alexa 488 and Alexa 568 conjugates (1:500, ThermoFisher) were used as secondary antibodies. Brain sections were imaged using an Axio Image D2 microscope (Zeiss) equipped with an AxioCam HRm camera.

### 8. ARNTL expression in cell-specific clusters using external single-nucleus RNA sequencing datasets

Processed publicly available single-nucleus RNA sequencing datasets of the mouse prefrontal cortex and hippocampus (Habib et al., 2017), the human dlPFC (Maitra et al., 2023), and human hippocampus (Ayhan et al., 2021) were downloaded, read into R (version 4.1.0) (https://www.R-project.org), and loaded as individual Seurat objects (version 4.0.4) (Hao et al., 2021). The clustering annotations for each dataset were also obtained. The DotPlot() function of Seurat was next used to assess the average expression of key regulators of the circadian rhythm, *ARNTL*, *DBP*, *CRY1*, *PER1* and *PER2,* in each cluster (broad and subclusters) of each dataset. For Maitra and colleagues’ (2023) dataset, which includes both neurotypical and depressed subjects, only nuclei from healthy individuals were included for analysis. Detailed information on the datasets used for analysis is presented in Supplementary Table 5.

### 9. ARNTL expression in region-specific clusters using an external spatial genomic dataset

The processed Seurat object of the human hippocampal Visium dataset generated by our group, which includes the clustering annotations, (GEO Accession: GSE248545) was loaded into R (version 4.1.0) (https://www.R-project.org). The DotPlot() function of Seurat (version 4.0.4) (Hao et al., 2021) was next used to assess the average expression of the core clock gene *ARNTL,* including the following canonical cell type markers: *SLC17A7* (excitatory neurons) *GAD1* (inhibitory neurons), *PROX1* (dentate lineage cells), *GFAP* (astrocytes), *TMEM119* (microglia), *MBP* (oligodendrocytes), *PDFGRA* (oligodendrocyte-precursor cells) and *RBFOX3* (neurons) in each BayesSpace (Zhao et al., 2021) cluster. Scaled values of expression levels are presented in the dot plot. Spatial plots were also created to spatially map the clusters and *ARNTL* expression in each section of the dataset at the spot level using the SpatialPlot() function of Seurat. The exact percentages and average expression levels of all markers in each cluster of the spatial transcriptomic dataset are presented in Supplementary Table 7.

### 10. ARNTL chromatin accessibility in cell-specific clusters using an external single-nucleus ATAC sequencing dataset

The snATAC-sequencing dataset was obtained from 40 neurotypical controls who died by natural causes or accidents. Cell type and cluster annotations as well as the open chromatin regions (OCRs) and cell type specific marker cCREs were obtained, as described (Chawla et al., 2023). Signac (Stuart et al., 2021) was used to calculate promoter accessibility for each gene based on the number of open chromatin fragments falling 2kbp upstream of gene TSS. For cis-regulatory activity of gene, peak fragments were integrated across entire gene body and signal was scaled bi-directionally using exponential decays from the gene TSS (extended upstream by 5 kb) and TTS, while accounting for neighboring gene boundaries (‘Gene Body Extended + Exponential Decay + Gene Boundary’) (Corces et al., 2020).

The most significant peak to gene links (Pearson’s r>0.5 & FDR<1e-4) across all cell types were identified using Pearson’s correlations between snATAC-sequencing accessibility at OCRs and matched snRNA-sequencing integrated gene expression, as described (Chawla et al., 2023; Corces et al., 2020). Homer (Heinz et al., 2010) (with default settings) was used to identify TF binding motifs enriched (BH corrected p<0.05) in peaks (n=9) which were positively correlated with ARNTL gene expression using peak to gene links (FDR<0.05). ARNTL TF binding sites in cell type OCRs were extracted as binary peak-by-motif match matrix (CIS-BP motifs), as described (Chawla et al., 2023; Corces et al., 2020). Finally, gene-ontology (GO) enrichments were performed using ClusterProfiler (Yu et al., 2012) and genome-wide annotations for humans (“org.Hs.eg.db”) with default settings (BH-corrected p<0.05).

### 11. Statistical analysis

Statistical analyses were performed using GraphPad Prism software (version 7). Normality and lognormality tests of variances were performed with D’Agostino & Pearson and Shapiro-Wilk tests. For comparisons between two groups, an unpaired two-sided Student’s *t*-test was used in the case of normally distributed data and a nonparametric test (Mann–Whitney *U* test) was applied in cases in which normality could not be assumed. Subjects with times of death between 6:00AM and 5:59PM were categorized in the “Day” group, and those with times of death between 6:00PM-5:59AM were categorized in the “Night” group, as previously done by our group (Belliveau et al., 2024). For comparisons between more than two groups, data were analyzed by a one-way ANOVA test in the case of normally distributed data, and we corrected for multiple comparisons using respectively Tukey’s. For correlation tests between dependent variables and covariates (age, PMI, pH and refrigeration delay), a two-sided Pearson’s correlation test was used in cases in which normality could be assumed and a Spearman’s correlation rank was performed in cases in which normality could not be assumed. A two-tailed approach was used for all tests and p-values < 0.05 were considered significant. Data is presented as mean ± SEM, unless specified otherwise.

## Acknowledgements

We would like to extend our deepest gratitude to the next of kin who consented to donating the brains of their loved ones. We also would like to thank the Douglas-Bell Canada Brain Bank staff and the Molecular and Cellular Microscopy Platform staff of the Douglas Research Centre (Montreal, Canada). This work was supported by a CIHR Project grant to NM. RR received postdoctoral fellowships from AMH and the FRQ-S.

## SUPPLEMENTARY FIGURE LEGENDS

**Supplementary Figure 1: *BMAL1* expression in *RBFOX3*-enriched regions of the human brain**

Representative images of *BMAL1* and *RBFOX3* expression in the adult human dlPFC and DG with DAPI staining, labeling nuclei, in blue. Scale bars = 1 mm.

**Supplementary Figure 2. Relationship between the percentage of *BMAL1*^+^ cells and age, PMI, pH, refrigeration delay, and time of death.**

(A) Pearson correlation between the percentage of *BMAL1*^+^ cells and age (r = −0.1760, p = 0.1010). (B) Pearson correlation between the percentage of *BMAL1*^+^ cells and PMI (r = −0.2600, p = 0.01444). (C) Pearson correlation between the percentage of *BMAL1*^+^ cells and pH (r = - 0.06644, p = 0.5386). (D) Pearson correlation between the percentage of *BMAL1*^+^ cells and refrigeration delay (r =-0.2142, p=0.04512). (E) Percentage of *BMAL1*^+^ cells in individuals with a time of death during daytime (Day) and in those with a time of death during nighttime (Night) (two-tailed Student t-test, p = 0.0032). Bar plot shows mean with SEM. (F) Percentage of *BMAL1*^+^ expressing *RBFOX3* or *P2RY12* in each brain region. Bar plot shows mean with SEM. (G) Percentage of *P2RY12*+ cells expressing *BMAL1* in each brain region Each dot corresponds to one region of interest (ROI) (from one staining replicate per subject). ** p ≤ 0.01 in two-tailed Student t-test.

**Supplementary Figure 3 Relationship between the percentage of *BMAL1*^+^ cells expressing *RBFOX3* and age, PMI, pH, refrigeration delay, and time of death.**

(A) Spearman correlation between the percentage of *BMAL1*^+^ cells expressing *RBFOX3* and age (r = 0.06710, p=0.5345). (B) Spearman correlation between the percentage of *BMAL1*^+^ cells expressing *RBFOX3* and PMI (r = −0.006675, p = 0.9508). (C) Spearman correlation between the percentage of *BMAL1*^+^ cells expressing *RBFOX3* and pH (r=0.07284, p = 0.5000). (D) Spearman correlation between the percentage of *BMAL1*^+^ cells expressing *RBFOX3* and refrigeration delay (r = 0.04991, p = 0.6442). (E) Percentage of *BMAL1*^+^ cells expressing *RBFOX3* in individuals with a time of death during daytime (Day) and in those with a time of death during nighttime (Night) (two-tailed Mann Whitney test, p = 0.8795). Bar plot shows mean with SEM. Each dot corresponds to one region of interest (ROI) (from one staining replicate per subject).

**Supplementary Figure 4. Relationship between the percentage of *BMAL1*^+^ cells expressing *P2RY12* and age, PMI, pH, refrigeration delay, and time of death.**

(A) Spearman correlation between the percentage of *BMAL1*^+^ cells expressing *P2RY12* and age (r=-0.1860, p=0.08271). (B) Spearman correlation between the percentage of *BMAL1*^+^ cells expressing *P2RY12* and PMI (r =0.05907, p = 0.5846). (C) Spearman correlation between the percentage of *BMAL1*^+^ cells expressing *P2RY12* and pH (r = 0.03381, p = 0.7545). (D) Spearman correlation between the percentage of *BMAL1*^+^ cells expressing *P2RY12* and refrigeration delay (r = −0.02181, p = 0.8402). (E) Percentage of *BMAL1*^+^ cells expressing *P2RY12* in individuals with a time of death during daytime (day group) and in those with a time of death during nighttime (night group) (two-tailed Mann Whitney test, p = 0.8721). Bar plot shows mean with SEM. Each dot corresponds to one region of interest (ROI) (from one staining replicate per subject).

**Supplementary Figure 5: Bmal1 protein expression in the mouse S1 barrel field, corpus callosum, striatum, and globus pallidus.**

(A) Representative confocal image of Bmal1 and Iba1 expression (with expanded insets at higher magnification) in the mouse S1 barrel field. Bmal1 and Iba1 do not co-localize. White arrows point to Iba^+^Bmal1^−^ cells. (B) Bmal1 and Iba1 expression (with expanded insets at higher magnification) in the mouse corpus callosum near the ventral hippocampal commissure. Lack of co-localization between Bmal1 and Iba1. White arrows point to Iba^+^Bmal1^−^ cells. (C) Bmal1 colocalizes with the astrocytic marker GFAP (with expanded insets at higher magnification) in the mouse corpus callosum near the lateral ventricle. White arrows point to GFAP^+^Bmal1^+^ cells. (D) Bmal1 and Iba1 expression (with expanded insets at higher magnification) in the mouse striatum and globus pallidus. Bmal1 and Iba1 do not co-localize. Scale bars = 50 µm.

**Supplementary Figure 6. ARNTL TF accessibility in a single-nucleus ATAC sequencing dataset.**

(A) TF binding sites enriched in microglial OCRs postiviely correlated wih ARNTL gene expression (Pearson’s correlation, FDR<0.05). (B) Violin Plots show TF motif accessibility for each cell type, ordered by average motif accessibility of ARNTL (BMAL1) in each cell type (Chawla et al., 2023). One-way ANOVA with Tukey HSD was performed for pairwise comparison between cell types. (C) Gene ontology enrichment (BH corrected p<0.05) showing functional pathways enriched for genes linked to ARNTL TF binding sites overlapping microglia OCRs (peak-to-gene linkages, r>0.45). Source data is presented is Supplementary Table 7.

**Supplementary Figure 7. Gene ontology enrichment (BH corrected p<0.05) showing functional pathways enriched for genes linked to ARNTL TF binding sites overlapping excitatory neuronal OCRs (peak-to-gene linkages, r>0.45).**

**Supplementary Figure 8. *BMAL2* expression in neuronal populations of the human dlPFC**

(A) Violin plot shows promoter (2kbp upstream of TSS) accessibility for ARNTL2 (BMAL2) gene, ordered by average expression of ARNTL in each cell type from Chawla et al. (2023). (B) Ridge plots show chromatin accessibility across cis-regulatory regions of ARNTL2 (5kbp upstream of TSS using an exponential decay model). (C) Multi-modal plots shows chromatin accessibility in each cell type (normalized by reads in TSS) at ARNTL2 gene locus (50kbp upstream and downstream of ARNTL2 TSS), followed by binarized single-cell coverage for 100 sampled cells from each cell type, and OCRs in each cell type in this locus. Finally, we show peak-to-gene linkages (Pearson’s r>0.5) between *ARNTL2* gene-expression and their regulatory peaks in this locus. (D) Scatter Plot shows R-squared value for correlations between ARNTL2 cis-regulatory activity (5kbp) and (E) ARNTL2 promoter accessibility, with age of each subject. (F) Representative confocal images of cells co-expressing *ARNTL* and *ARNTL2* in the human dlPFC grey matter of a sudden-death individuals. Scale bar = 50 µm. (G) Average percentage of *BMAL1*^+^ cells expressing *BMAL2*, *P2RY12* or both markers in the human dlPFC (n=3). For each section, four regions of interest were counted. Each dot corresponds to one region of interest (from one staining replicate per subject). Bar plot shows mean with SEM.

**Supplementary Figure 9. Relationship between ARNTL promoter accessibility, cis-regulatory activity, and gene activity and age.**

Scatter Plot shows R-squared value for correlations between ARNTL (BMAL1) promoter accessibility (A), cis-regulatory activity (5kbp) (B), and gene activity (C), and age of each subject.

## SUPPLEMENTARY TABLE LEGENDS

**Supplementary Table 1. Subject information for RNAscope and CD11^+^ microglia isolation experiments.**

**Supplementary Table 2. Source data for percentages of *BMAL1*^+^ cells expressing *P2RY12* and *RBFOX3* in different brain regions (from RNAscope/FISH experiments).**

**Supplementary Table 3. List of detected proteins and their abundance from LC-MS proteomics analysis in the mouse brain and post-mortem human vmPFC grey matter.**

**Supplementary Table 4. Source data for percentages of *BMAL1*^+^ cells expressing *ALDH1L1* or *RBFOX3* in the dlPFC.**

**Supplementary Table 5. Source data for the percentages of nuclei expressing *Arntl/ARNTL* in cell-type specific clusters from Habib et al. (2017), Ayhan et al. (2021) and Maitra et al. (2023).**

**Supplementary Table 6. Source data for the percentages of spots expressing *ARNTL* in region-type specific clusters from Simard et al. (2024).**

**Supplementary Table 7. Microglia OCRs showing significant peak-to-gene correlation with *BMAL1* expression (used for TF motif enrichment).**

**Supplementary Table 8. Gene ontology enrichment analysis showing functional pathways enriched for genes linked to ARNTL TF binding sites overlapping microglia OCRs.**

## Notes

### Competing Interest Statement

The authors have declared no competing interest.

